# Evaluating brain parcellations using the distance controlled boundary coefficient

**DOI:** 10.1101/2021.05.11.443151

**Authors:** Da Zhi, Maedbh King, Carlos R. Hernandez-Castillo, Jörn Diedrichsen

## Abstract

One important approach to human brain mapping is to define a set of distinct regions that can be linked to unique functions. Numerous brain parcellations have been proposed, using cytoarchitectonic, structural or functional Magnetic Resonance Imaging (fMRI) data. The intrinsic smoothness of brain data, however, poses a problem for current methods seeking to compare different parcellations. For example, criteria that simply compare within-parcel to between-parcel similarity provide even random parcellations with a high value. Furthermore, the evaluation is biased by the spatial scale of the parcellation. To address this problem, we propose the Distance Controlled Boundary Coefficient (DCBC), an unbiased criterion to evaluate discrete parcellations. We employ this new criterion to evaluate existing parcellations of the human neocortex in their power to predict functional boundaries for an fMRI data set with many different tasks, as well as for resting-state data. We find that common anatomical parcellations do not perform better than chance, suggesting that task-based functional boundaries do not align well with sulcal landmarks. Parcellations based on resting-state fMRI data perform well; in some cases, as well as a parcellation defined on the evaluation data itself. Finally, multi-modal parcellations that combine functional and anatomical criteria perform substantially worse than those based on functional data alone, indicating that functionally homogeneous regions often span major anatomical landmarks. Overall, the DCBC advances the field of functional brain mapping by providing an unbiased metric that compares the predictive ability of different brain parcellations to define brain regions that are functionally maximally distinct.

## 1. Introduction

Neuroscience has a long history of subdividing the human brain into different regions based on differences in histology (Brodmann, 1909). It is commonly understood that brain function arises through the interactions of regions that are structurally and/or functionally distinct (Felleman and Van Essen, 1991, Eickhoff et al., 2018). While early parcellations of the human brain were based on the cytoarchitectonic organisation of the neocortex (Brodmann, 1909, Zilles et al., 2002, Talairach, 1988), the advent of neuroimaging allowed an in-vivo assessment of brain organisation. In recent years, many parcellations based on task-evoked (Yeo et al., 2015) and resting functional magnetic imaging resonance (fMRI) data (Eickhoff et al., 2015, Arslan et al., 2018, Eickhoff et al., 2018) have been published, along with multi-modal parcellations that also incorporate structural and cytoarchitectonic information (Glasser and Van Essen, 2011, Fan et al., 2016).

In the empirical study of brain function, parcellations play an important practical role. They are commonly used to define the regions of interest (ROIs) to summarize functional and anatomical data, or to define the nodes for subsequent connectivity analysis (Sporns, 2011). The main function of parcellation is to reduce complexity of the statistical analysis, as the brain-wide data can be summarized with a smaller number of values, each reflecting measurements from a region with high homogeneity. Additionally, widely-accepted parcellations aid the direct comparison between studies (Arslan et al., 2018).

Despite the importance of brain parcellations in human neuroscience research, there is no commonly accepted evaluation criterion to compare different parcellations. The obvious reason for this is that different parcellations are generated with different goals in mind. Specially, some parcellations aim to define regions that have a common anatomical characteristic (Desikan et al., 2006, Fischl et al., 2004), a shared connectivity fingerprint (Yeo et al., 2011, Gordon et al., 2016, Power et al., 2011), or a homogeneous task-activation profile (Yeo et al., 2015). As such, brain parcellations can be evaluated based on different types of data (Arslan et al., 2018).

Universally, however, any parcellation should aim to define regions such that the functional profiles (whether anatomical measures, connectivity patterns, or task activation) of two brain locations in the same region should be maximally similar to each other, whereas two brain locations in different regions should be maximally different. Thus, brain parcellation can be viewed as a clustering problem. As a result, standard machine learning methods to evaluate clustering solutions have been applied to brain parcellation. Two such examples are the measure of global Homogeneity (Gordon et al., 2016, Craddock et al., 2012) and the Silhouette coefficient (Rousseeuw, 1987).

However, these two evaluation criteria have the common problem in that they do not account for the spatial nature of the underlying data. In the case of the human neocortex, the functional correlation between two nodes on the cortical surface depends on their distance, with nearby nodes showing a higher similarity compared to far away ones. This causes even random, but spatially contiguous, parcellations to achieve relatively high global Homogeneity or Silhouette coefficient. To establish whether a parcellation identifies any real functional boundaries at all, Monte-Carlo simulations using random parcellations are therefore necessary (Arslan et al., 2018). To complicate matters further, both global Homogeneity and Silhouette coefficient tend to be higher for finer parcellations. This makes it difficult to compare between two parcellations with different spatial resolutions.

In this paper, we address this problem by proposing a novel evaluation criterion, the Distance-Controlled Boundary Coefficient (DCBC). As the Silhouette coefficient, it compares within-parcel and between-parcel correlations of the functional profiles. However, the DCBC takes into account the spatial smoothness of the data by only comparing pairs of locations with the same distance on the cortical surface. As we will show, the expected value of the DCBC for a random parcellation is zero. Thus, no simulations with random parcellations are necessary to establish a baseline measurement; we can directly test the DCBC against zero. We also show that this baseline value is invariant to the number of parcels in the random parcellation. This enables us to use the DCBC to directly compare parcellations of different spatial scales.

We then use the DCBC to evaluate a set of common parcellations of the human neocortex (Yeo et al., 2011, 2015, Gordon et al., 2016, Power et al., 2011, Glasser et al., 2016, Schaefer et al., 2018, Fan et al., 2016, Baldassano et al., 2015, Shen et al., 2013, Arslan et al., 2015, Tzourio-Mazoyer et al., 2002, Desikan et al., 2006, Fischl et al., 2004). We performed this evaluation using both a task-based and a resting-state fMRI data set. For the task-based data set, we used the comprehensive Multi-Domain Task Battery (MDTB) (King et al., 2019), which contains functional contrasts across many cognitive domains measured in the same participants. A python toolbox for the efficient computation of the DCBC, as well as a surface-based version of the MDTB data set are publicly available to download.

## 2. Methods

### 2.1. Overview

The DCBC compares the correlation between two brain locations within a parcel to the correlation between two brain locations across a boundary between parcels. Importantly, this comparison is only performed for pairs of brain locations that are separated by the same spatial distance. The calculation of the DCBC proceeds in four steps. First, we require a data set that provides a rich characterization of each brain location. This data set defines the *functional profile* for each brain location. While the DCBC can be applied to any high-dimensional data, such as multi-modal anatomical data, we focus here on task-based fMRI data (the MDTB data set (King et al., 2019), which provides 34 activity estimates across a range of motor, cognitive and social tasks) and resting-state fMRI data (acquired in the Human Connectome Project, HCP, (Van Essen et al., 2013)). Secondly, we need a measure of spatial distance between two brain locations, either defined on the cortical surface, or for subcortical structures, in the volume. Based on these distances, all location pairs are subdivided into a set of spatial bins. The within-parcel and between-parcel correlation is then computed for each spatial bin separately. In the last step, the results are integrated across spatial bins, using an adaptive weighting scheme. To validate the method, we employed random parcellations of the human neocortex using a range of spatial resolutions, as well as sets of smooth artificial functional data sets.

### 2.2. Evaluation Data

#### 2.2.1. Task-based dataset (MDTB)

To define the functional profiles for the evaluation, we first used the publicly available MDTB data set (King et al., 2019), which contains a wide range of tasks, quantifying processes required for motor, cognitive, and social function. Each of the 24 participants (16 females, 8 males, mean age=23.8) was scanned four times for 80-minutes, while performing either task set A or B (17 tasks for each, 9 tasks in common). Task set A was performed in the first two sessions, task set B in the last two sessions. A total of approximately 5.3 h of functional data per participant was collected.

In each imaging run, every task was performed once for 35 s, starting with a 5 s instruction period, followed by a 30 s period of continuous task performance. The task battery included motor (finger tapping, sequence production), working memory (2-back task, math), language (verb generation, reading), social (theory of mind, action observation), executive control (no-go, stroop), attention (visual search), emotion (facial expression, pleasant/ unpleasant pictures), spatial (mental rotations), introspection tasks (spatial and motor imagery), movie-based tasks (cartoon, nature, landscapes), and rest (fixation) (King et al., 2019).

All fMRI data were acquired on a 3T Siemens Prisma at Western University. The imaging parameters were as follows: repetition time = 1 s; field-of-view = 20.8 cm; phase encoding direction P to A; 48 slices; 3 mm thickness; in-plane resolution 2.5×2.5 *mm*^2^. For anatomical localization and normalization, a 5 min high-resolution scan of the whole brain was acquired (see (King et al., 2019) for more details).

Data pre-processing was carried out using tools from SPM12 (www.fil.ion.ucl.ac.uk/spm/doc/spm12_manual.pdf), as well as custom-written scripts written in MATLAB. For all participants, an anatomical image (T1-weighted MPRAGE, 1mm isotropic resolution) was acquired in the first scanning session. Functional data were realigned for head motion within each session, and for different head positions across sessions using the six-parameter rigid body transformation. The mean functional image was then co-registered to the anatomical image and this transformation was applied to all functional images. No smoothing or group normalization was applied.

The anatomical image of each of the 24 subjects was processed by standard recon-all pipeline of the freesurfer software (version 5.0) (Fischl, 2012), including brain extraction, white and pial surfaces generation, inflation, and spherical alignment to the new symmetric fsLR-32K template (Van Essen et al., 2012). Individual surfaces were then re-sampled into this standard grid. This resampling led to surfaces with 32,492 vertices that are shared both across participants and across left and right hemisphere.

A General Linear Model (GLM) was fitted to the time series data of each voxel for each imaging run. Each task was modeled as a 30s regressor and all the preceding 5s instructions were modeled as separate regressors. The regression weights (betas) were estimated for each run independently and then averaged across the 16 runs for each task set.

To combine the activity estimates across the two task sets, we used the mean of the shared tasks as a common reference point. We subtracted this pattern from the average beta estimates for each task set separately, and then concatenated the two vectors of activity estimates. The average beta weights were then divided by the square root of the average mean-square-residual from the first-level GLM to obtain z-scores for each voxel. The resulting functional profiles consisted of 34 pre-whitened activity estimates (set A = 17; set B = 17) for each voxel. Finally, we subtracted the overall mean across all tasks from the functional profile of each voxel.

The functional profiles were then mapped to each individual cortical surface by averaging the value from voxels along the connecting line between the pial and white-gray matter surface, using 5 equally spaced locations between the two surfaces.

#### 2.2.2. Resting-state dataset (HCP)

The second data set used in this study was the resting-state fMRI (rs-fMRI) data from the “unrelated 100” subjects (54 female, 46 male adults, aged from 22 to 35), which was made publicly available in the Human Connectome Project (HCP) S1200 release (Van Essen et al., 2013). The rs-fMRI scans for each subject were collected in two sessions held on different days, including a total four runs of approximately 15 minutes each. During the scans, the subjects were asked to fixate a white cross-hair on a dark background.

The HCP resting-state fMRI time series were acquired using 3T Siemens “Connectome Skyra” scanner with 2 × 2 × 2 mm spatial resolution and a TR of approximately 0.7 s. For more details of the data acquisition parameters, see Smith et al. (2013), and Uğurbil et al. (2013).

All data were pre-processed using the HCP minimal processing pipeline (Glasser et al., 2013), including structural registration, correction for spatial distortion, head motion, cortical surface mapping, and functional artefact removal (Smith et al., 2013, Glasser et al., 2013). For each rs-fMRI run, this resulted in 1200 time points for each of the 32k vertices of the standard fsLR-32K template (Van Essen et al., 2012) per hemisphere. To generate the functional profiles for the HCP data set, we concatenated all 4 runs after mean-centering.

### 2.3. Existing evaluation criteria for brain parcellations

Given that brain parcellation can be viewed as a clustering problem, two common methods used to evaluate the resultant parcels are the global Homogeneity (Craddock et al., 2012, Gordon et al., 2016), and the Silhouette coefficient (Rousseeuw, 1987). Homogeneity is defined as the average similarity across all pairs of vertices within a parcel. As the similarity measure of two vertices, we used the Pearson’s correlation between functional profiles. The global Homogeneity is then simply the average with-parcel correlation across all parcels, with higher homogeneity suggesting a better parcellation.

Another popular evaluation metric for brain parcellations is the Silhouette coefficient (Rousseeuw, 1987), which compares the average dissimilarity (defined as 1-R, where R represents Pearson’s correlation between functional profiles) from one vertex to all other vertices in the same parcel (*w*_*i*_), to the average dissimilarity from the same vertex to all the vertices that assigned to neighbouring parcels (*b*_*i*_) (Yeo et al., 2011, Arslan et al., 2018). For a given a parcellation {*P*_1_,*P*_2_,…,*P*_*k*_}, *w*_*i*_ and *b*_*i*_ can be defined as:

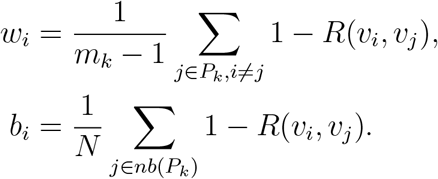

where *m*_*k*_ indicates the number of vertices that within the parcel *P*_*k*_. *N* is the total number of vertices in all neighbouring parcels and *nb*(*P*_*k*_) represents all neighbouring parcels of *P*_*k*_.

For each cortical vertex *v*_*i*_, the Silhouette coefficient is defined as:

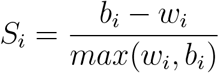

Based on this definition, the Silhouette coefficient for each vertex ranges from -1 to 1, 1 indicates that there is a perfect correlation within each parcel (*r* = 1) and on average, not correlations across parcels (*r* = 0). As we will see, both of these measures are biased by the intrinsic smoothness of the functional profiles on the cortical surface.

### 2.4. Measuring spatial distance

To account for the intrinsic smoothness of the data, we require a measure of spatial distance between any pair of brain locations. For subcortical structures, we have used the Euclidean distance between pairs of voxels (King et al., 2019). For the neocortex, however, we ideally would like to use the geodesic distance between vertices on the cortical surface. As an approximation to this distance, we used Dijkstra’s algorithm (Dijkstra et al., 1959) to estimate the shortest paths between each pair of vertices on each individual cortical surface. For this computation we used the mid-cortical layer which is the average of the pial and white-gray matter surface. For computational and memory efficiency we only considered distances up to maximum of 50mm. Inter-vertex distances were then averaged across individuals and hemispheres. This resulted in a matrix that indicates the average cortical distance between nearby brain locations for the atlas brain surface.

### 2.5. Distance Controlled Boundary Coefficient (DCBC)

#### 2.5.1. The problem of spatial smoothness

The problem with global Homogeneity and Silhouette coefficient is that they do not take account that function tends to vary in a smooth fashion across the cortical surface. For instance, if we compute the correlation of vertex pairs across the cortex using task-evoked functional profiles (King et al., 2019) for an individual participant, we can clearly see that the correlation falls off with the spatial distance between vertices (See Figure 1a). Note that this smoothness is not an artifact of the data processing; except for motion realignment and mapping onto the surface, no smoothing was applied to the data. Thus, the dependence on spatial distance reflects the intrinsic smoothness of functional specialization on the cortex.

**Figure 1:**
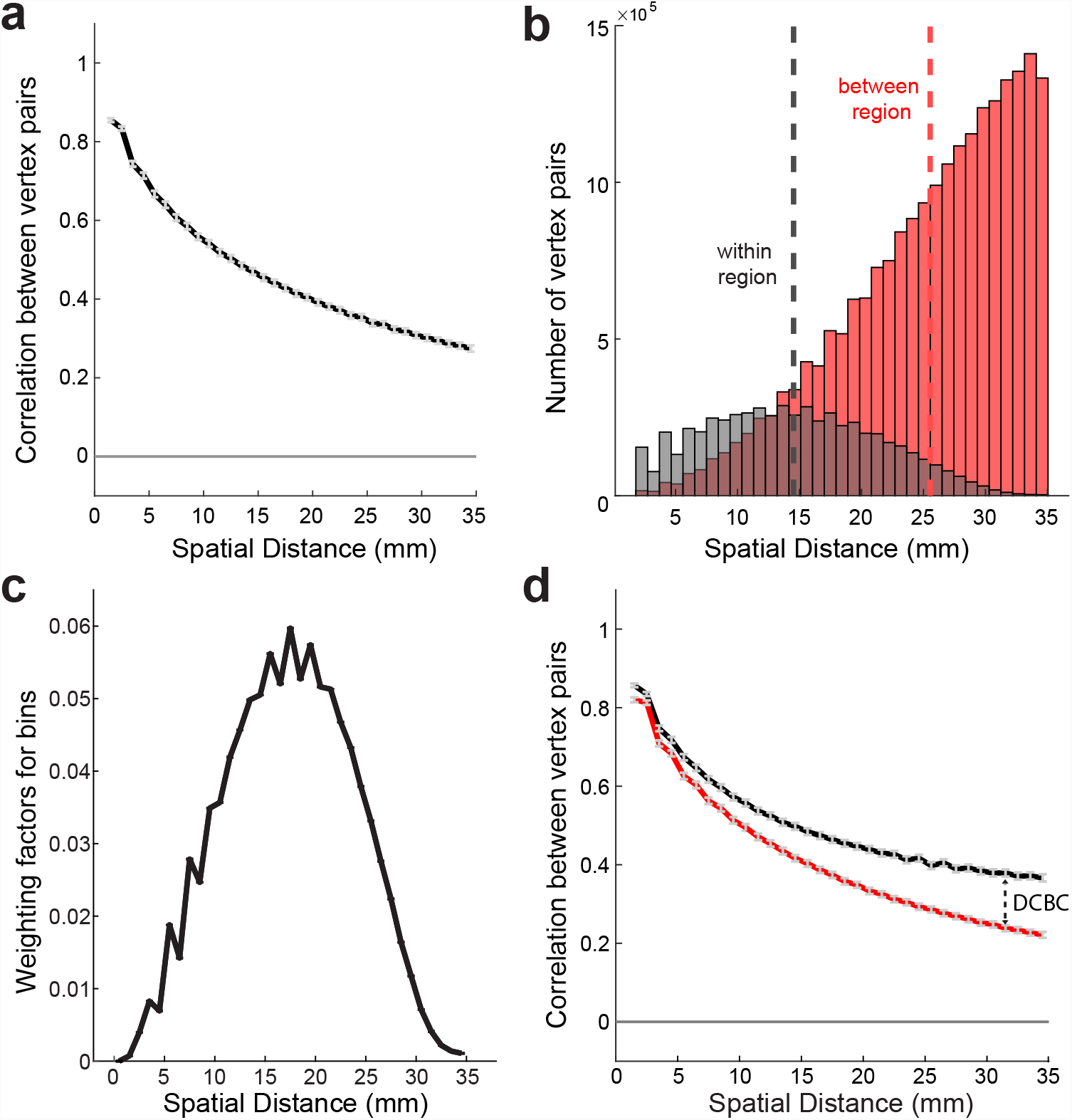
Distance controlled boundary coefficient. (a) Correlation between task-evoked functional profiles (see methods) of pairs of surface vertices as a function of their spatial distance; (b) Histogram of the number of within and between vertex pairs as a function of spatial distance for a random Icosahedron 162 parcellation. (c) Weighting factor across different bins for the Icosahedron 162 parcellation and binning shown in b. (d) The correlations for within- (black) and between- region (red) vertex pairs as a function of the spatial distance (for Yeo 17 parcellation). The DCBC is defined by the weighted average distance between the two curves.

**Figure 2:**
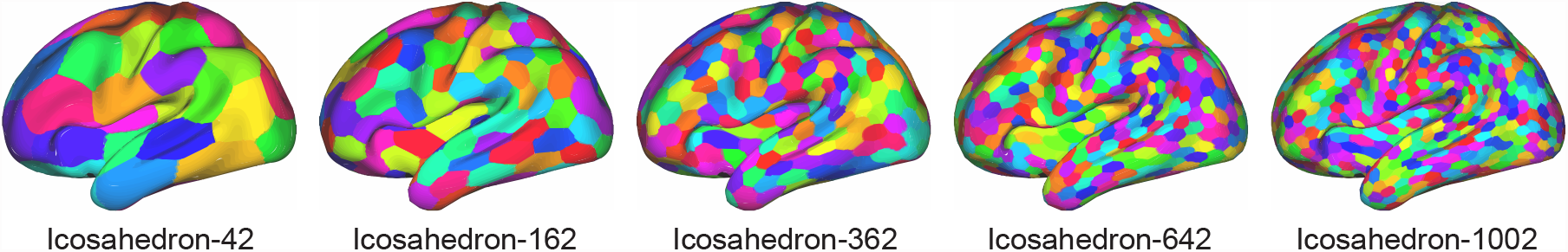
Random cortical parcellations with different number of parcels.

For the global Homogeneity measure, this property favors parcellations with small parcels, as only close-by vertex pairs will be within the same parcel. Similarly, the spatial smoothness also biases the Silhouette coefficient, as the spatial distance for within-parcel pairs is on average smaller than that for between-parcel pairs. For example in random parcellation Icosahedron 162 (Fig. 1b), the average spatial distance of within-parcel pairs is 14.5 mm.

Even if we limit the comparison to vertex pairs from spatially adjacent parcels, as is common practice in the evaluation of brain parcellations, the between-parcel pairs have a substantially larger average distance (25.5mm). This discrepancy results in a higher average correlation of functional profiles for within-parcel pairs compared to between-parcel pairs.

We therefore propose to only compare vertex pairs with a similar spatial distance. For this purpose, the DCBC method bins all vertex pairs according to their spatial distance, and then compares the correlation for within- and between-pairs within each bin. One important practical decision is the choice of bin size, a question that we address in the results section. For our neocortical data, a bin size of 1 mm appears to be adequate.

#### 2.5.2. Averaging across bins

Parcellations can be compared by investigating the difference in within- and between-parcels as a function of the spatial distance (see King et al. (2019), Fig. 3,4). However, for many applications we would like a single evaluation criterion for each parcellation, which necessitates the averaging across a range of spatial distances. This raises the question of what range of spatial distances to consider, and how to weight the distances within that range. A rational solution to this problem is to find the weighting that, for any given parcellation, provides us with the best estimate of the average difference between within- and between-parcel correlations, assuming that this differences is constant across the desired range of distances. The variance of the estimate of the correlation difference (*d*_*i*_) for bin *i* can be approximated by assuming the independence of the different vertex pairs. In this case, the variance of the estimate depends on the number of within- (*n*_*w,i*_) and between-parcel vertex pairs (*n*_*b,i*_) in each spatial bin:

**Figure 3:**
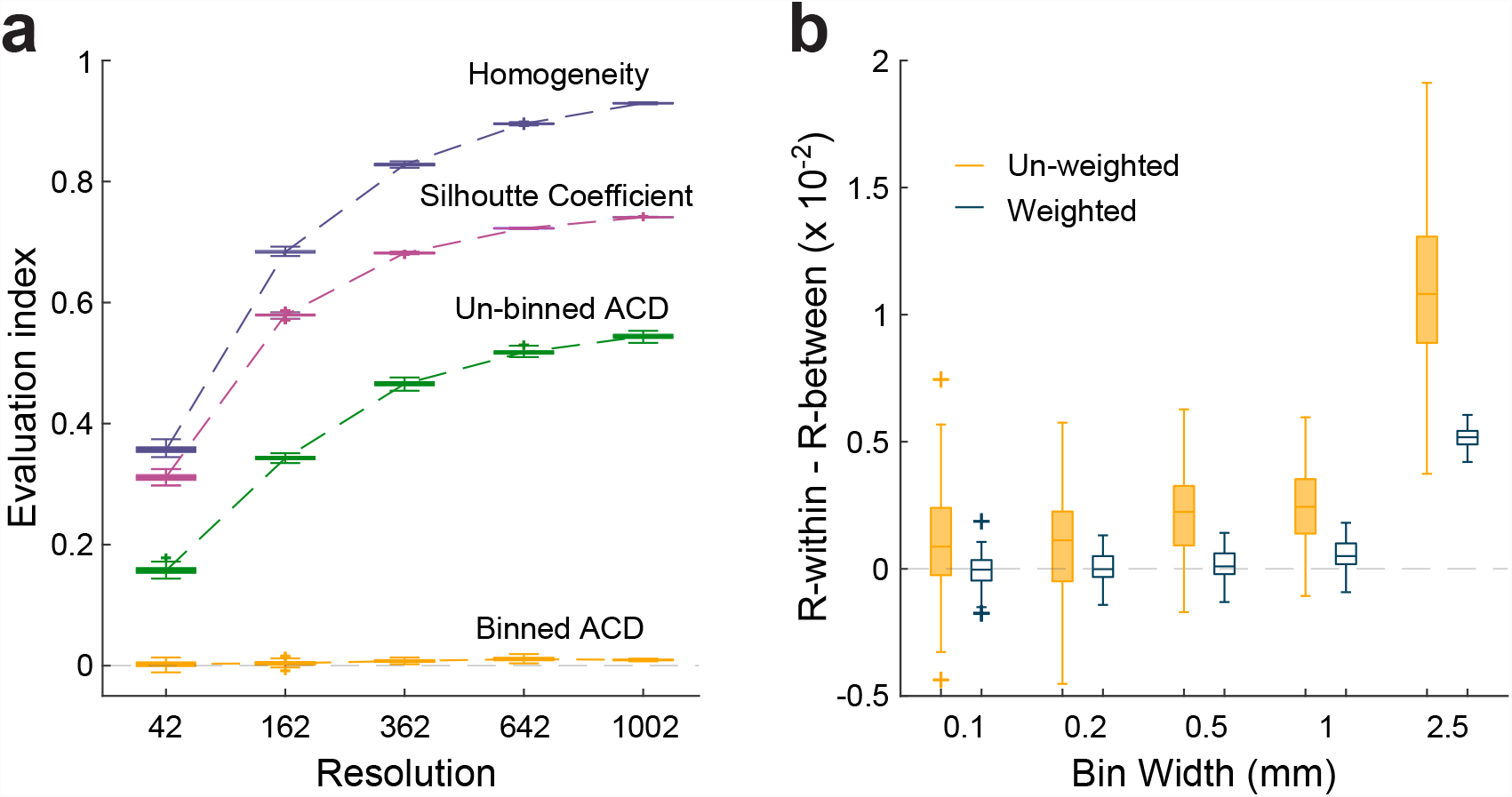
(a) Results of four evaluation methods for the random functional maps simulation in different resolutions: Homogeneity, Silhouette Coefficient, the un-binned average correlation difference (ACD) of within-parcel and between-parcel, and the binned (bin width = 2.5 mm) ACD of within-parcel and between-parcel; (b) The weighted and un-weighted DCBC evaluation results for 100 random functional maps in different bin widths of random parcellations Icosahedron 642.

**Figure 4:**
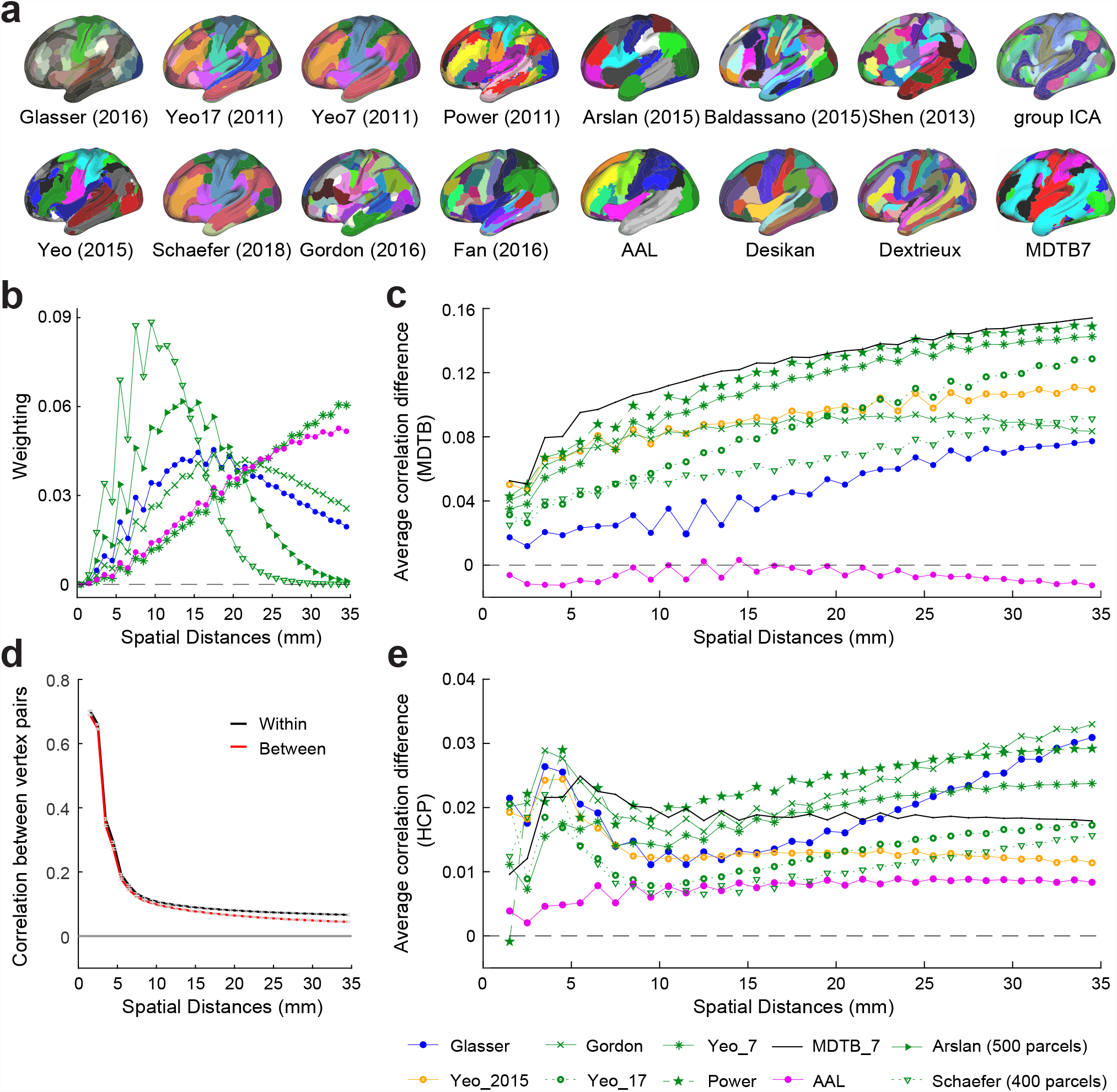
Evaluation on the real data sets. (a) The left hemisphere of 15 commonly used cortical parcellations and the MDTB cortical parcellation with 7 regions; (b) weighting factors of selected parcellations with different resolutions within range of 0 to 35mm; (c) the correlation difference as a function of the spatial distance for selected parcellations evaluated on the task-based data set; (d) the correlation for within- and between-parcel vertex pairs as a function of the spatial distance (for Yeo 17 parcellation) evaluated on the resting-state dataset; (e) the average correlation difference as a function of the spatial distances for selected parcellations evaluated on the resting-state data set.

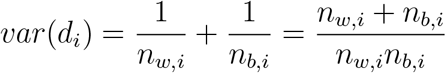

For averaging, we define a weight that is proportional to the precision (inverse of the variance) of each estimator:

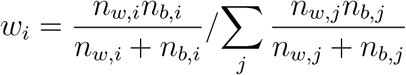

For example, figure 1c shows the weighting factor for each spatial bin of Icosahedron 162 random parcellation using a 1 mm bin width. The DCBC is then the weighted average of the correlation difference across bins.

### 2.6. Random Parcellations

To evaluate the DCBC for parcellations that on average do not align with real functional boundaries, we generated a set of random parcellations. If our method successfully controls for the spatial smoothness of the functional profiles, the average DCBC for such random parcellations should be zero, i.e there should be no difference between within- and between-parcel correlations. To test this claim for parcellations at different spatial scales, we used a regular hexagonal parcellations of a sphere (Icosahedron) with 42, 162, 362, 642, and 1002 parcels. To generate random alignment of this parcellation with the data, we rotated each map randomly around the x, y, and z axis. We repeated this process 100 times to obtain 100 random parcellations for each spatial scale.

### 2.7. Random Functional Maps

Real data may have functional boundaries that are correlated across participants. As a consequence, some random parcellations will still by chance align with these boundaries and yield systematically positive DCBC values; and other random parcellations will misalign with the real functional boundaries and yield systematically negative values. To test the DCBC on functional maps without a systematic boundary structure across participants, we also randomly generated 100 cortical functional maps with 34 task conditions. These maps then were used in the analysis shown in Fig. 3. We drew independent normally distributed values for every condition and vertex for the fsLR-32k template (Van Essen et al., 2012), and then smoothed these maps on the cortical surface using `-metric-smooth` function provided by Connectome Workbench software (Marcus et al., 2011). The smoothing kernel was set to 12 centimeters, yielding a similar spatial autocorrelation function as in our real data.

### 2.8. Evaluation of commonly-used group parcellations

We then evaluated a set of commonly used group parcellations of the human neocortex (Table 1). The majority of the parcellations considered here are based on resting-state functional connectivity (rs-FC) data. The fMRI data is recorded at rest, and the covariance or correlations between the time series of different brain locations is submitted to a clustering or dimensionality reduction approach. Parcellations can also be based on task-evoked activation maps. For example Yeo et al. (2015) derived a parcellation from 10,449 experiment contrasts across 83 behavioural tasks. The anatomical parcellations considered here are based on the macro-anatomical folding structure of the human neocortex, following the major sulci and gyri of the human brain. Finally, we also considered 2 multi-modal parcellations, which combined rs-FC and anatomical criteria.

**Table 1:**
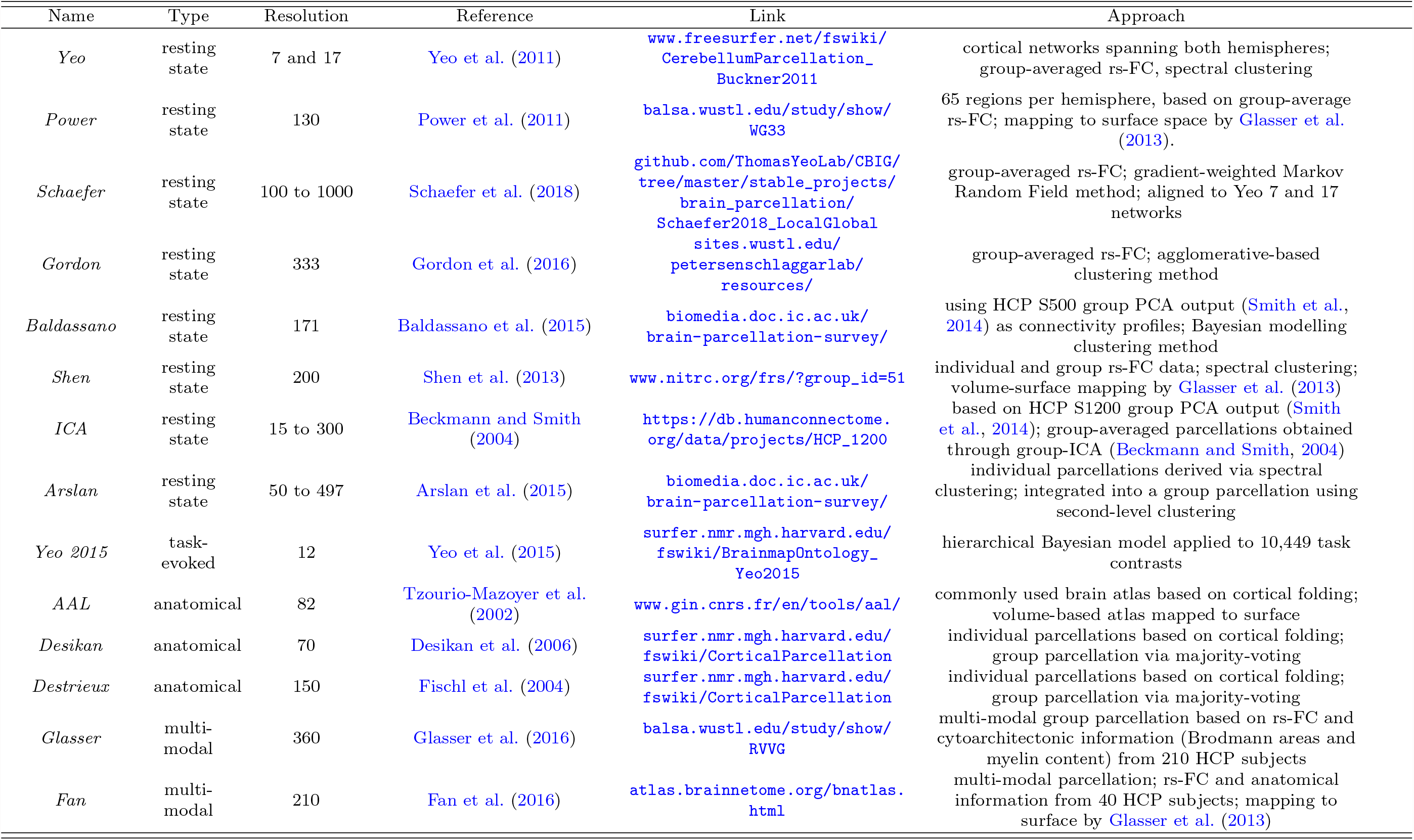
Commonly used group-level cortical parcellations. rs-FC: resting state functional connectivity (fMRI).

Within each of these categories, parcellations also differ in the approach used for generation. For instance, several parcellations were obtained based on a two-level approach, where subject-level parcellations are derived in a first step, and then integrated across subjects in a second step using clustering or majority voting. Other parcellations are directly derived by clustering group-averaged data (Arslan et al., 2018).

Because the DCBC evaluation considers only vertex pairs up to a specific spatial distance on the cortical surface, the evaluation is conducted separately for the left and right hemisphere. For many parcellations, the parcels are separated for the two hemispheres. For example, Gordon et al. (2016) used 161 and 172 distinct regions for the left and right hemisphere respectively, totaling 333 regions. Other parcellations use bi-hemispheric parcels. As a consequence the 7 and 17 regions in Yeo et al. (2011), were effectively evaluated as 14 and 34 parcels.

Note that three group-level parcellations (Fan et al., 2016, Shen et al., 2013, Tzourio-Mazoyer et al., 2002) were only available in volume space. These parcellations were mapped to HCP standard fsLR-32k cortical surface using the volume-to-surface pipeline described in Van Essen et al. (2012) and Arslan et al. (2018). All parcellations in this study are available as a collection in fsLR-32k surface space at (Zhi and Diedrichsen, 2021).

### 2.9. Parcellation based on the evaluation data

To estimate how well a group parcellation could theoretically subdivide the neocortex into functionally distinct regions, we derived a parcellation from the MDTB dataset. We estimated 12 cortical parcellations with 14 to 1000 parcels, using group-averaged MTDB functional profiles. For evaluation on the MDTB data, this parcellation has the unfair advantage that the individuals used in the evaluation is also contained within the training set, providing an upper-bound estimate of the noise ceiling (Nili et al., 2014). To estimate a lower bound of the noise ceiling, we used a leave-one-out cross validation approach: We derived a group parcellation from the averaged data from 23 participants, and then evaluated it on the remaining subject. We then averaged the DCBC across the 24 different parcellations.

To derive the MDTB group parcellation we used spectral clustering. We first down-sampled the surface data from 32K vertices to 4002 vertices. Then we performed spectral clustering on the affinity matrix between the vertices of the down-sampled map. After clustering, we then restored the map to the original resolution of the surface. The lower resolution ensured that the resulting parcellations were spatially contiguous. We consider the MDTB group parcellation as a potential lower bound (see Results 3.4 and Discussion) of how well a group parcellation can perform on the MDTB data set.

## 3. Results

### 3.1. Binning reduces the bias introduced by spatial smoothness

Existing evaluation methods for brain parcellations have the problem of being biased by the natural smoothness of functional brain maps. To demonstrate this effect, we first tested various evaluation methods using random functional maps and random parcellations of different spatial scales. As can be seen in Fig. 3a, both the Homogeneity and Silhouette coefficient show highly significant positive values, even for these random maps. Furthermore, the values for both methods increase when the parcellation increases in spatial resolution (i.e. have more and smaller parcels). This makes direct comparisons of different parcellations in different spatial scales difficult, and necessitates the use of randomisation analyses for each parcellation to determine the baseline value expected by random chance (Arslan et al., 2018).

A similar problem can also be seen when using the difference between the correlations of within- and between-parcel pairs of vertices (un-binned average correlation difference, ACD). This is caused by the tendency that vertices that are closer together show higher functional correlations (Fig. 1a), combined with the fact that within-parcel vertex pairs are on average closer to each other than between-parcel pairs (Fig. 1b). To remove this bias, the DCBC calculation involves the binning of vertex pairs according to their spatial distance. We then calculate the difference between the average correlation between within- and between-parcel pairs within each spatial bin, thereby only comparing vertex pairs of matched spatial distance.

To ascertain that this approach provides an approximately unbiased and stable evaluation criterion, we first applied the suggested technique on the simulated functional data. As can be seen (Fig. 3a, binned ACD), even using a relatively coarse spatial binning of 2.5 mm, this approach nearly removes all bias caused by the spatial smoothness. For the finest parcellation, an Icosahedron with 1002 regions, the binned difference between correlations (0.009) was approximately 60 times lower than the mean of the difference calculated in each bin (0.544). This shows that the binning approach dramatically reduces the bias caused by spatial smoothness.

### 3.2. Adaptive weighted averaging reduces variance and bias

After binning, we often want to integrate the results across bins to arrive at a single evaluation criterion. This can be done by simply averaging the differences in correlation across bins. However, this approach leads to a summary measure with high variability (Fig. 3b). This is caused by the fact all bins have equal influence on the average, even though some bins contain very few vertex pairs. This can be addressed by taking the number of within- and between-parcel pairs in each bin into account in an adaptive weighting scheme (see methods). Indeed, the standard deviation of the weighted DCBC in the simulation is 2.8 times lower than for the un-weighted version for 1 mm bins, and 8.1 times lower for the 2.5 mm bins. Furthermore, the weighted DCBC also shows smaller residual bias than the unweighted DCBC.

### 3.3. Choosing an appropriate bin width

An important practical issue in the DCBC calculation is to choose a specific bin width. This choice is subject to a fundamental trade-off. If the bin width is too wide, the DCBC may still be biased as a result of the spatial smoothness of the functional profiles. This is because within each bin, the average spatial distance for within-parcel pairs is still slightly smaller than for the between-parcel pairs, inducing a systematic difference between the correlations within each spatial bin. Even though this bias is small, it can remain highly significant when tested across the 100 simulations presented in Fig. 3b for a bin width of 2.5mm. Choosing a smaller bin width reduces this bias. For bins of size 0.1 mm and 0.2 mm, the same 100 simulated data sets no longer show a significant difference against zero (p = 0.327 and 0.202, respectively).

Choosing a very small bin width, however, also comes with some disadvantages. First, the computational cost of the DCBC calculation increases linearly with the increasing number of bins. More importantly, if a very small bin is chosen, it can result in noisier DCBC estimate, as very few vertex pairs will fall within each bin. In the extreme case, there would either be no within- or between- vertex pair in a bin, such that the bin could not be used for the difference calculation. For neocortical data projected to the fsLR-32k template (Van Essen et al., 2012) a bin width of 1mm appears to offer a good balance between bias, accuracy and computational speed.

### 3.4. DCBC evaluation for real data

Using a task-based data set (MDTB) and a resting-state data set (HCP), we evaluated 15 commonly used group-level cortical parcellations (Fig. 4a, Table 1). These parcellations relied either on anatomical criteria (cortical folding), task-evoked activation, or functional resting-state connectivity. Two multi-modal parcellations (Glasser et al., 2016) relied on a combination of anatomical and functional features. Each of the parcellations was evaluated per hemisphere and the global DCBC of a subject was then averaged across hemispheres.

For the MDTB data set, the difference between the within-parcel and between-parcel correlations across range of spatial distances (0-35mm) is shown in Fig. 4c. While the difference increased with increasing distance, the relative ordering of the parcellations was relatively consistent: Independent of the exact spatial distance considered, the Power and the MDTB correlation appear to outperform the other parcellations.

For the HCP data set (Fig. 4e), the difference between the within- and between-parcel correlation was substantially smaller. This is mostly caused by the fact that the raw Pearson’s correlation of the time series (Fig. 4d) were lower than the correlations for the MDTB data set (Fig. 1a,d). The correlations for the rs-fMRI data also fell off more rapidly with increasing distance, reaching values of *r* < 0.1 for distances over 8mm (Fig. 4d). The lower DCBC values for this data set, therefore, are partly due the fact that correlations between full fMRI time series are usually lower than correlations between task-based activity estimates. For the HCP data set the difference in correlations were relatively stable across the range of spatial distances considered (< 35*mm*).

To obtain a minimum-variance estimate of the correlation difference when averaging across spatial distances, we weighted the difference curves with the parcellation-specific weighting function (Fig. 4b). This procedure is certainly justified if the differences between correlation curves is stable across spatial distances. For parcellation where the differences between within- and between-parcel correlations vary with distance, small biases may arise. For example we may expect for the MDTB data set could give a small advantage to coarser parcellations. We will return to this issue in the discussion.

### 3.5. Resting-state group parcellations predict task-based functional boundaries

We then calculated the averaged weighted DCBC across all parcellation (Fig. 5). The first insight is that the nine parcellations that are based solely on functional resting state data (Yeo et al., 2011, Power et al., 2011, Gordon et al., 2016, Arslan et al., 2015, Baldassano et al., 2015, Shen et al., 2013, Schaefer et al., 2018, Smith et al., 2014) predicted the functional boundaries in the task-based data set substantially better than chance (Fig. 5a). For example, the within- and between-parcel correlations for the Yeo 17 parcellation (Fig. 1d) differed by approximately 0.1 across spatial bins, reflected in an average DCBC value of 0.1020 (SE=0.0053) across the 24 subjects. Other resting-state parcellations also showed clear differences between the within- and between-parcel correlations, especially Power 2011 (DCBC=0.1334, SE=0.0085), Yeo 7 (DCBC =0.1271, SE=0.0073), and Gordon 2016 (DCBC=0.0876, SE=0.0047). This finding confirms that resting-state group parcellations generally predict task-relevant functional boundaries significantly better than chance (Tavor et al., 2016).

**Figure 5:**
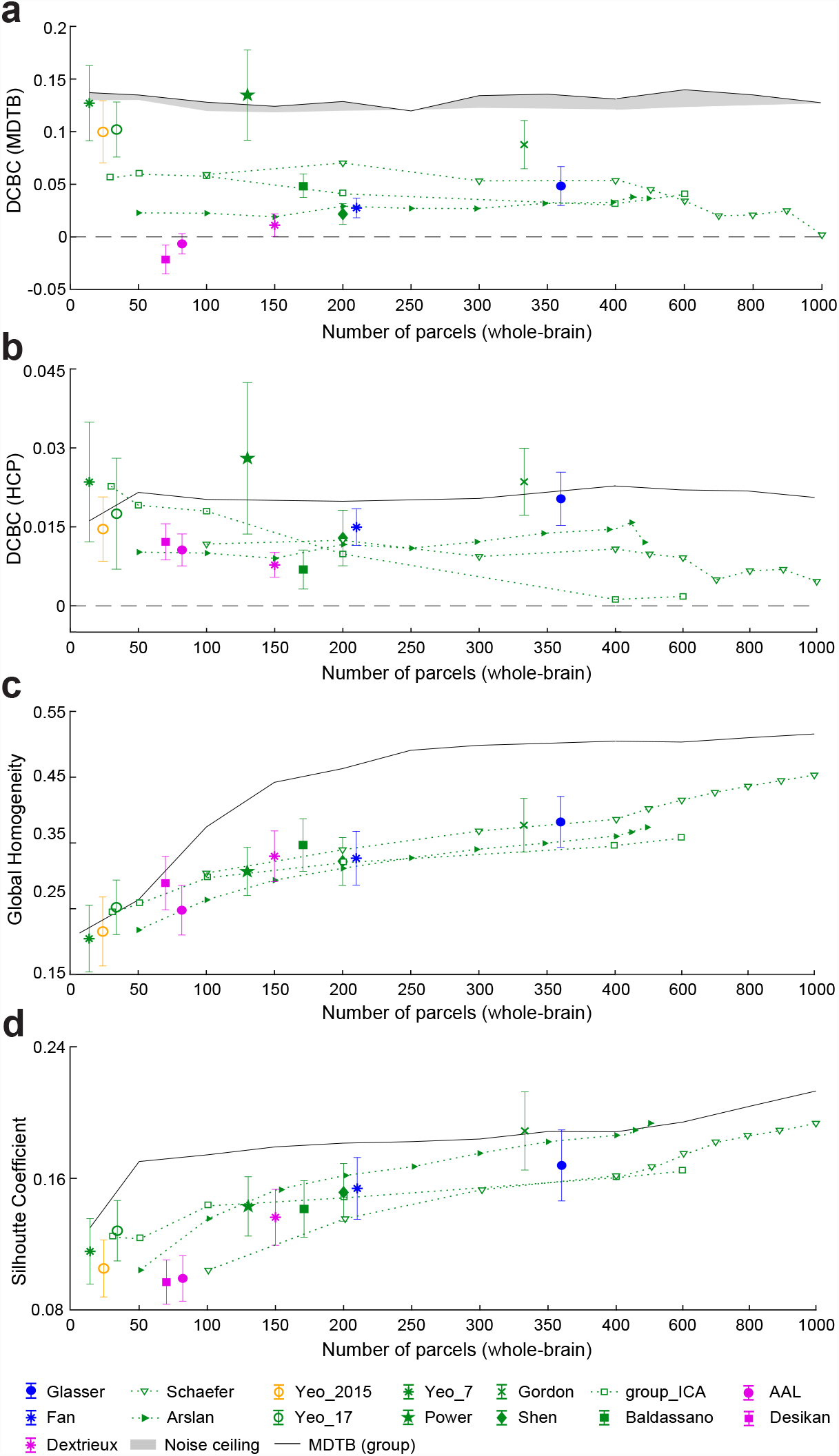
Evaluation of existing group parcellations. (a) The DCBC evaluation of the existing parcellations on the task-based dataset. The evaluation of the MDTB parcellations is indicated by the gray area (noise ceiling). The upper and lower bounds are the non-crossvalidated and cross-validated evaluations respectively; (b) the DCBC evaluation of the existing parcellations on HCP resting-state dataset; (c) the global homogeneity measures for all parcellations using the MDTB task-based dataset. Error bars indicate the standard deviation of DCBC across the 24 evaluation subjects. (d) The Silhouette Coefficient of the parcellations using the MDTB task-based dataset.

When evaluating these resting-state parcellations on resting state data (see Fig. 5b), we obtain consistent results. Even though the overall DCBC was substantially lower than for the task-based data, the best-performing parcellations were based on resting-state data, including the Yeo 7 (DCBC=0.0213, SE=0.0021), Power (DCBC=0.0261, SE=0.0025), and Gordon (DCBC=0.0236, SE=0.0018) parcellations.

### 3.6. Comparison to parcellations derived from the evaluation data set

How well do these group-based resting-state parcellations predict task-based functional boundaries, relative to what would be possible? Given the inter-individual variability of boundaries, and the fact that even individual boundaries are not perfectly sharp, there is an upper limit to the highest achievable DCBC on our evaluation data set. To obtain an idea of this “noise ceiling” (Nili et al., 2014), we derived a set of clustering solutions from the MDTB data itself (see methods, Fig. 4a), spanning the range from 14 to 1000 parcels.

While we cannot determine the noise ceiling directly, we can obtain a lower and upper performance estimate. For the lower estimate, we derived the parcellation on 23 of the participants, and evaluated it on the remaining, left-out participant. For the upper estimate, we over-fitted the data by deriving and evaluating the parcellation on all 24 participants. The gap between these two performance curves indicates how much of the performance advantage of the MDTB parcellation is due to the over-fitting to the particular set of subjects.

As expected, the MDTB-based parcellations (gray area in Fig. 5a) generally outperformed other group parcellations on this data set. Nonetheless, some existing resting-state parcellations showed performance very close or even slightly higher than the MDTB parcellation (Yeo et al., 2011, Power et al., 2011).

When evaluated the task-based MDTB parcellations on the resting-state data (Fig. 5b), it performed remarkably well, and was only outperformed by 3 resting-state parcellations. This again demonstrates the consistency of functional boundaries across task- and resting-state data.

### 3.7. Anatomical parcellations do not predict task-based functional boundaries

We then evaluated 3 commonly used anatomical group parcellations of the human neocortex: The Desikan parcellation (Desikan et al., 2006), the Dextrieux atlas (Fischl et al., 2004), and the Automated Anatomical Labelling (AAL) atlas (Tzourio-Mazoyer et al., 2002). On the task-based data (Fig. 5a), the averaged correlation between any vertex pairs within a predefined anatomical parcel was not much greater than the correlation between vertex pairs across anatomical boundaries, resulting in very low DCBC values (Dextrieux: DCBC=0.0112, SE=0.0022; AAL: DCBC=-0.0066, SE=0.002). The Desikan parcellation (DCBC=-0.0215, SE=0.0028) even showed significantly negative DCBC values across the 24 subjects. These parcellations, based on the macroanatomical folding structure of the neocortex, therefore define boundaries that are systematically misaligned with the functional subdivisions in task-evoked activity profiles.

A similar pattern emerged when anatomical parcellation were evaluated on resting-state data. All anatomical parcellations showed relatively low performance (average DCBC = 0.0066). In contrast to the task-based evaluation, the DCBC of all three parcellations was significantly positive, when tested against zero (all *t*_23_ *>* 9.4466, *p* < 2.21 × 10^−9^), implying that they aligned with the boundaries of the resting-state networks slightly better than chance.

### 3.8. Multi-modal parcellations do not perform better than resting-state parcellations

We also applied DCBC evaluation to two multi-modal parcellations (Glasser et al., 2016, Fan et al., 2016) to determine whether combining anatomical and functional data is superior to unimodal parcellations. The Glasser parcellation had a higher DCBC score (DCBC=0.0483, SE=0.0038) as compared to the Fan parcellation (DCBC=0.0275, SE=0.0019). However, both were lower than the average DCBC across the resting-state parcellations (0.0766). It therefore appears that the combination of multiple modalities does not systematically lead to a better prediction of task-relevant function boundaries than parcellations that are derived on resting-state data alone.

### 3.9. Comparison across different spatial resolutions

For simulated random functional maps, we have shown that the expected value of the DCBC is zero, no matter how fine the parcellation (Fig. 3b). In contrast, the value of the global Homogeneity and Silhouette coefficient increases for finer parcellations even for random maps (Fig. 3a).

This bias can also be observed for real parcellations. The value of the global Homogeneity (Fig. 5c) and Silhouette coefficient (Fig. 5d) when calculated on the task-based evaluation data set clearly increases for finer parcellations, whereas there is no strong relationship between the DCBC and the number of parcels (Fig. 5a and b).

In this context, the set of Schaefer 2018 parcellations (Schaefer et al., 2018) is especially interesting, as it provides a nested set of subdivisions with an increasing number of parcels, all aligned with Yeo 7 or 17 networks (we use the one aligned with Yeo 7 networks in the experiment). To statistically evaluate the change in evaluation metric with parcel size, we conducted a repeated-measures analysis of variance (ANOVA) across the 10 Schaefer parcellations, ranging from 100 to 1000 parcels. As expected, both the Homogeneity (*F*_9,207_ = 1730.6, *p* = 1.55 × 10^−189^) and the Silhouette coefficient (*F*_9,207_ = 667.6, *p* = 1.11 × 10^−147^) clearly showed significant increases for an increasing number of parcels. Given that such increases were also found for random functional maps and parcellations, it is not clear whether the finer parcellations identified functional boundaries better, the same, or worse than coarser parcellations.

In contrast, the unbiased DCBC shows that the Schaefer parcellation reaches the best performance around 200 parcels, and then slowly declines for finer parcellations (Fig. 5a). The ANOVA showed a significant change with number of parcels (*F*_9,207_ = 189.4576, *p* = 8.19 × 10^−95^). Indeed, for the finest parcellation (1000 parcels), performance did not differ significantly from chance (*t*_23_ = 1.0253, *p* = 0.3159). One possible reason for this is that when defining more than 200 functional parcels, the new boundaries do not consistently predict discontinuities in the functional organisation at the group level anymore.

In summary, the application of the novel DCBC criterion to known cortical parcellations allowed for the following conclusions: (1) anatomical parcellations based on cortical folding do not align with functional boundaries in the neocortex; (2) resting-state parcellations predict task-relevant functional boundaries, outperforming other types of cortical parcellations; (3) multi-modal parcellations did not improve performance as compared to pure resting-state parcellations.

### 3.10. Open-source toolbox/data support evaluation

The code for DCBC evaluation is published as an open-source toolbox written in Python at (Zhi and Diedrichsen, 2021). The package also contains the pre-processed contrast maps for all task conditions of the MDTB data set (n=24 subjects), sampled to the standard fsLR-32k template.

## 4. Discussion

In this study, we introduce a novel evaluation criterion for brain parcellations, the Distance Controlled Boundary Coefficient (DCBC). The method takes into account the spatial smoothness of the data by controlling the distance of the vertex pairs when comparing within- and between-parcel correlations. We used an earlier form of this approach for volume-based data (using the Euclidean distance instead of a surface-based distance) to evaluate cerebellar parcellations (King et al., 2019). Here, we further improve the approach by adaptively weighting over the spatial distance bins, resulting in a global measure with low bias and variance. Our evaluation on simulated smooth data shows that the new criterion overcomes the size- and shape-dependent bias of other homogeneity-type evaluation criteria (Craddock et al., 2012, Rousseeuw, 1987). The advantage of the DCBC is twofold: 1) it enables a direct comparison between group brain parcellations that have different spatial resolutions, and 2) it provides a direct test of how well a parcellation subdivides the brain into homogeneous regions than expected by chance.

One important caveat is that DCBC only removes the bias completely if the difference between within- and between correlations is stable across spatial distances. This is because different parcellations use different weighting across spatial distances (Fig. 4b). There are a number of practical solutions to ameliorate this problem. First, by choosing a maximal distance for vertex distances (here 35*mm*) the evaluation is constrained to be relatively local in all cases. While future users of the method may want to choose a different range of spatial distances, we believe that 35mm provides a good compromise for cortical parcellations. Secondly, it is always useful to plot the DCBC as a function of the spatial distance before averaging (see King et al. (2019), Fig. 4c,e) to investigate whether different parcellations may behave differently across the distances considered. Finally, if the DCBC varies substantially across spatial distances, one could use a common, averaged weighting for all parcellations, or simply decide on a more specific set of spatial distances. Nonetheless, the biases from differential weighting were relatively small for our evaluation data sets - and the DCBC successfully removed the main biasing influence of parcel size (Fig. 5a vs. c,d).

We used the DCBC to evaluate a range of existing cortical surface-based parcellations in their ability to predict functional boundaries on task-based and resting-state data. We found that the parcellations derived from resting-state fMRI data largely succeed in predicting task-evoked activity boundaries, replicating earlier work (Tavor et al., 2016, King et al., 2019). These results demonstrate again the practical utility of resting state data in identifying brain networks that work together during active task performance. Even though the correlation structure across the cortex does clearly change in a task-dependent fashion (Hasson et al., 2009, Salehi et al., 2020a,b), our results emphasize the existence of a basic scaffold that determines functional specialization across a wide range of tasks, as well as during rest. In the opposite direction, parcellations derived from a rich task-based battery (MDTB) also achieved relatively high DCBC values when evaluated on rs-fMRI data, further confirming that structure of neural fluctuations during rest aligns with co-activation across tasks.

In contrast, anatomical parcellations (Desikan et al., 2006, Fischl et al., 2004, Tzourio-Mazoyer et al., 2002) did not perform better than chance to predict functional boundaries in the task-based data, and only slightly better than random for rs-fMRI data. The Desikan parcellation even showed a negative DCBC score on the MDTB dataset. This finding corroborates previous work that shows a misalignment between macroanatomical folding structure and functional boundaries in the neocortex (Arslan et al., 2018) and the cerebellum (King et al., 2019). An inspection of the differences between functional and anatomical parcellations (Fig. 4a) suggest an explanation of why this may be the case. Cortical motor areas, for example, are subdivided in all anatomical parcellations along the central sulcus, which separates the primary motor cortex (M1) from primary somatosensory cortex (S1). In this case, the macro-anatomical folding roughly aligns with the cyto-architectonic boundaries between the two regions (Brodmann, 1909, Amunts and Zilles, 2015, Fischl et al., 2008). In contrast, functional parcellations typically separate the motor regions along a ventral-dorsal axis, that is, into foot, hand and mouth regions. Within each body zone, these parcellations leave M1, S1, and premotor regions in the same parcel, likely reflecting the high functional correlations between regions responsible for the control of a body part. Similar observations can be made in the visual system - with functional parcellations separating areas associated with foveal and peripheral vision, rather than subdividing known cytoarchitectonic regions (V1, V2, V3). This anti-correlation of functional and anatomical boundaries partly explains why the Desikan atlas showed a significantly negative DCBC.

It is therefore also unsurprising that multi-modal parcellations that combine functional and anatomical criteria did not outperform the pure resting-state parcellations. For example, Glasser et al. (2016) used resting-state connectivity, intra-cortical myelin signal, and cortical folding, thereby improving alignment with cytoarchitectonically defined areas. The inclusion of anatomical information likely also led to the division of functionally correlated brain regions. This does not imply that cytoarchitectonic parcellations of the neocortex are of lesser value. Instead, our finding simply illustrates that the goal of isolating anatomically consistently organised regions is unrelated to, and in some cases conflicts with, the goal of defining brain regions with homogeneous functional profiles.

Therefore, the evaluation results in our study would have a different look if we used anatomical data rather than task-evoked activity profiles as an evaluation data set. It is worth stressing, however, that the DCBC as a method to control for the influence of spatial smoothness can be used with any suitable data set. For instance, anatomical data, such as myelin measures (Tozer et al., 2005) or anatomical connectivity fingerprints derived from diffusion data (Behrens et al., 2003, Johansen-Berg et al., 2004) could be used to evaluate how well parcellations isolate anatomically homogeneous regions.

In the current study we focused on task-evoked and resting-state fMRI data as evaluation targets. While the two datasets led to a similar pattern of results when comparing parcellations, the overall DCBC values for rs-fMRI data were substantially lower that the DCBC values for task-based fMRI data. This is likely due to the lower signal-to-noise level for fMRI time-series, as compared to task activity estimates, which are averaged over 16 runs. Note that the high correlation for vertex pairs with small spatial distances (*<*5mm) are likely driven by the interpolation across neighboring voxels (motion realignment, surface mapping, minimal smoothing) inherent in both pre-processing pipelines. One advantage of evaluating parcellations on task-activity data is the obvious face validity of the results: If the goal of the the brain parcellation is to define regions with a homogeneous response across a wide range of tasks and mental states, then it should be best to evaluate the parcellation that way, rather than relying on the possibly more restricted mental states during rest.

A possible extension of the current work is to develop a parcellation algorithm that explicitly optimizes the DCBC. Given the nature of the DCBC, such an algorithm would have to be a local, rather than a global parcellation algorithm (Schaefer et al., 2018), such as a region-growing algorithm proposed in Gordon et al. (2016) or Salehi et al. (2020a). The algorithm to be proposed would very likely improve the DCBC beyond what was achieved by spectral clustering, which does not consider the spatial arrangement of the vertices.

Even more substantial improvement in the quality of the parcellations can be expected when moving from a group to an individual parcellation. Recent results have shown that the inter-individual differences in the exact spatial location of functional boundaries are substantial (Salehi et al., 2020b, King et al., 2019). Of course, individual parcellations can suffer from having too little individual data to reliably establish the parcellation. Optimal ways of fusing group and individual-level data (Buckner et al., 2013, Kong et al., 2019), which also makes parcels comparable across subjects (Salehi et al., 2020a) clearly provides a promising future addition to the neuroimaging toolkit. In these efforts, the DCBC can provide a bias-free and reliable evaluation criterion that can be calculated without computationally expensive simulation studies.

When developing, using, and evaluating brain parcellations, it is of course important to consider the much more fundamental issue of whether this form of representation (Bijsterbosch et al., 2020) is a valid description of brain organisation. In our mind, it remains an open question to what degree segmenting the brain into discrete regions with hard boundaries (Van Essen and Glasser, 2018) is a useful description at all, or to what degree this functional organisation is better explained through a set of smooth gradients (Tononi et al., 1994, Dohmatob et al., 2021, Huntenburg et al., 2018, Guell et al., 2018). Either way, we believe that the DCBC evaluation provides an useful tool to advance this debate. If brain functions only varied in smooth gradients across the cortical surface, the DCBC should not be systematically above zero, at least not when evaluated on a novel set of tasks. However, most resting-state parcellations identified boundaries that also aligned with more rapid changes in the active functional response. Thus, as for the human cerebellum (King et al., 2019), this demonstrates the existence of task-invariant functional boundaries on the cortical surface. On the other hand, not all boundaries are equally strong, and not all boundaries are equally stable across tasks. The ability of DCBC to evaluate individual boundaries, as done in King et al. (2019), therefore provides an important tool to evaluate both functional segregation, as well as continuous functional integration (Eickhoff et al., 2018) in a region-specific way.

A Python-based software toolbox for the evaluation of surface-based parcellations on the MDTB activity maps is made publicly available at (Zhi and Diedrichsen, 2021). The toolbox is also designed to allow users to evaluate parcellations on other types of data. We hope that the new evaluation criterion will support and facilitate researchers in understanding the functional compartmentalization of the human brain.

## 5. Acknowledgements

This study was supported by a Discovery Grant from the Natural Sciences and Engineering Research Council of Canada (NSERC, RGPIN-2016-04890), and a project grant from the Canadian Institutes of Health Research (CIHR, PJT 159520), both to J.D. Additional funding came from the Canada First Research Excellence Fund (BrainsCAN) to Western University.

